# Effector-dependent response deterioration by stochastic transformations reveals mixed reference frames for decisions

**DOI:** 10.1101/193235

**Authors:** T. Scott Murdison, Dominic Standage, Philippe Lefèvre, Gunnar Blohm

## Abstract

Recent psychophysical and modeling studies have revealed that sensorimotor reference frame transformations (RFTs) add variability to motor output by decreasing the fidelity of sensory signals. How RFT stochasticity affects the sensory input underlying perceptual decisions, if at all, is unknown. To investigate this, we asked participants to perform a simple two-alternative motion direction discrimination task under varying conditions of head roll and/or stimulus rotation while responding either with a saccade or button press, allowing us to attribute behavioral effects to eye-, head- and shoulder-centered reference frames. We observed a rotation-induced, increase in reaction time and decrease in accuracy, indicating a degradation of motion evidence commensurate with a decrease in motion strength. Inter-participant differences in performance were best explained by a continuum of eye-head-shoulder representations of accumulated decision evidence, with eye- and shoulder-centered preferences during saccades and button presses, respectively. We argue that perceptual decision making and stochastic RFTs are inseparable, consistent with electrophysiological recordings in neural areas thought to be encoding sensorimotor signals for perceptual decisions. Furthermore, transformational stochasticity appears to be a generalized phenomenon, applicable throughout the perceptual and motor systems. We show for the first time that, by simply rolling one’s head, perceptual decision making is impaired in a way that is captured by stochastic RFTs.

**Significance statement:** When exploring our environment, we typically maintain upright head orientations, often even despite increased energy expenditure. One possible explanation for this apparently suboptimal behavior might come from the finding that sensorimotor transformations, required for generating geometrically-correct behavior, add signal- dependent variability (stochasticity) to perception and action. Here, we explore the functional interaction of stochastic transformations and perceptual decisions by rolling the head and/or stimulus during a motion direction discrimination task. We find that, during visuomotor rotations, perceptual decisions are significantly impaired in both speed and accuracy in a way that is captured by stochastic transformations. Thus, our findings suggest that keeping one’s head aligned with gravity is in fact ideal for making perceptual judgments about our environment.

## Introduction

We typically maintain upright head and eye orientations with respect to the horizon (Pozzo et al., 1990; Dunbar et al., 2004, 2008), despite potentially increased energy expenditure. For example, during hunting (Land, 2014), flight (Altshuler et al., 2015) or motorcycle racing it would be more energy efficient to align the head with the inertial vector. Minimizing vertical disparity has been suggested as one reason for this behavior (Misslisch et al., 2001; Schreiber et al., 2001). A potential complementary reason could come from the recent finding that reference frame transformations (RFTs) are stochastic (Alikhanian et al., 2015), as is apparent in both perception (Schlicht and Schrater, 2007; Burns et al., 2011) and motor planning (Sober and Sabes, 2003, 2005; McGuire and Sabes, 2009; Burns and Blohm, 2010). If the encoding of evidence is similarly degraded by stochastic transformations, then maintaining specific head orientations while making visuomotor decisions could be optimal for the signal’s preservation, despite requiring energy expenditure.

Bounded accumulator models account for a wealth of behavioral data from perceptual decision tasks under the premise that noisy evidence for the alternatives is accumulated until it reaches a criterion bound (Smith and Ratcliff, 2004; Bogacz et al., 2006). Under this framework, stochastic RFTs could influence choice behavior in predictable ways. One possibility is that RFTs can degrade the encoding of evidence by lowering its signal-to-noise ratio. In this case, the behavioral outcome should be commensurate with increasing task difficulty, resulting in increased reaction times (RTs) and decreased accuracy (percent correct).

The goal of this study was to determine the influence of stochastic RFTs on perceptual decision making. To do so, participants were asked to perform a 2AFC motion direction discrimination task either under non-rotated (control) conditions or under several different head roll or rotated stimulus conditions (Figure 1). In a blocked design, they were also instructed to indicate their decision regarding the left or right direction of coherent motion with either a saccade or a button press. Because eye movements are executed in head-centered coordinates and, when the arm is stationary, button presses occur in shoulder-centered coordinates, this paradigm allowed us to perform well-established psychometric and chronometric analyses while also allowing us to test the effects of eye-, head- and shoulder-reference frames on choice behavior.

**Figure 1:**
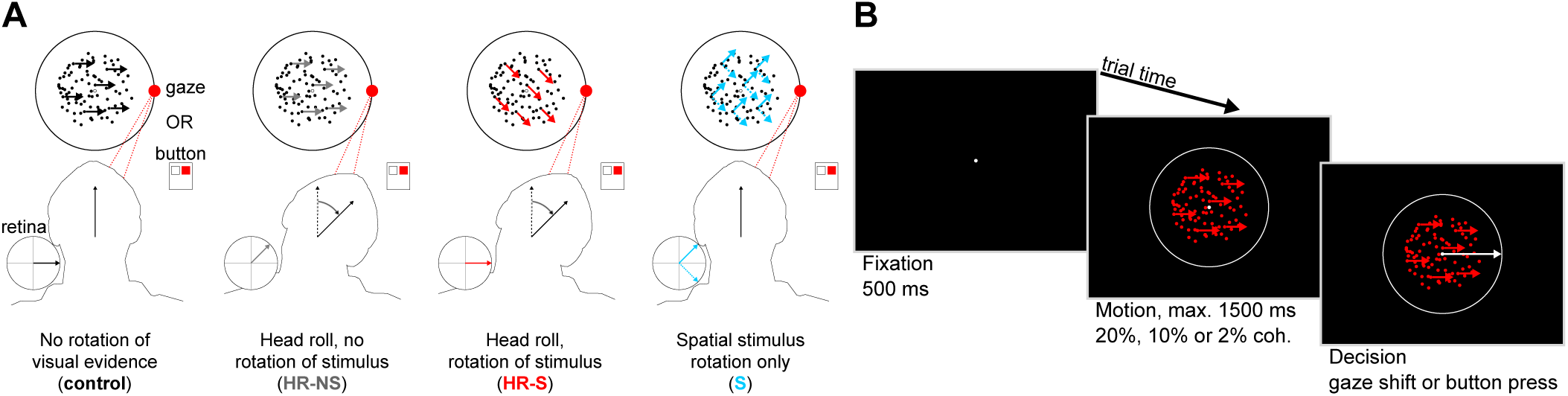
Task and paradigm. (A) Participants performed the task under one of eight conditions – four for each response type (saccade or button), organized in a block design. These were combinations of head and/or congruent screen rotations, giving rise to visual motion that was separable across eye, head and shoulder (screen) reference frames. (B) Each trial consisted of a fixation (500 ms), motion (up to 1500 ms) and decision epoch. Participants were instructed to determine the direction (left or right) of coherently moving dots randomly chosen at 20%, 10% or 2% coherence and make their decision using either a horizontal saccade or a button press as quickly and accurately as possible.

## Materials and methods

### Experimental paradigm

To test how reference frame transformations affect perceptual decisions, we developed an experimental paradigm with distinct conditions consisting of (1) rotations of the visual stimulus, (2) rotations of the head and (3) changes to the response type (saccade or button press). These conditions allowed us to comprehensively investigate the influence of different reference frame transformations on the decision process based on the coding frame of the motion evidence and transformation of that evidence into a reference frame appropriate for the motor response. These conditions are illustrated in Figure 1A.

We determined participants’ baseline decision making performance using a control condition in which participants’ heads remained upright (0° roll) and the axis of coherent motion remained along the horizontal (0°) screen-centered axis. Thus, comparing our other experimental conditions to this one provided the effects directly resulting from adding new requirements to the transformation (Figure 1A, first column). For each response type, the rotational conditions were rolling the participants’ heads towards a shoulder (about 45°), without rotation of the on-screen stimulus (head roll – no stimulus rotation, HR-NS, Figure 1A, second column); head roll with 45° rotation of the on-screen stimulus (head roll – stimulus rotation, HR-S, Figure 1A, third column); 45° rotation of only the on-screen stimulus (S, Figure 1A, fourth column).

### Participants

In total, 12 participants (age 20-32 years, 8 male) were recruited for two experiments after informed consent was obtained. Eleven of 12 participants were right-handed and 11 of 12 participants were naïve as to the purpose of each experiment (main and control). Each experiment had seven participants, and two participants performed both main and control experiments. Participants in the main experiment were between the ages of 22 and 32 years (5 male) and all were right-handed. Participants in the control experiment were between the ages of 20 and 26 years (4 male) and six of seven were right-handed. All participants had normal or corrected-to-normal vision and did not have any known neurological, oculomotor, or visual disorders. All procedures were approved by the Queen’s University Ethics Committee in compliance with the Declaration of Helsinki.

### Apparatus

Participants sat in complete darkness 50 cm in front of a 36 cm x 27 cm Dell UltraScan P991 CRT monitor (Dell, Round Rock, TX). Participants’ heads rested on a chin rest that allowed for head roll in the frontoparallel plane. With their heads in an upright position on the chin rest, the interocular midpoint was aligned to the frontoparallel fixation position on the screen. The visual stimulus was displayed on the screen (120 Hz refresh rate) using the ViSaGe Visual Stimulus Generator with VSG Toolbox for Matlab (Cambridge Research Systems, Rochester, UK). Movements of both eyes were recorded at 400 Hz using a Chronos head-mounted 3D video eye tracker (Chronos Vision, Berlin, Germany) that was stabilized to the head using a bite bar. Head movements were recorded at 400 Hz using an Optotrak Certus system (Northern Digital, Waterloo, Ontario, Canada) with three infrared diode markers placed on the Chronos helmet. For consistency across camera positions, these helmet markers were calibrated with respect to an external orthonormal axis defined by a set of three orthogonal diodes located either on the wall behind the participant or on the side of the CRT monitor. Screen brightness and contrast settings were adjusted so that participants could not see the edges of the monitor screen in complete darkness, even after 0.5 h dark adaptation.

### Procedure

The visual stimulus consisted of a centered array of white circular dots (0.1° diameter) arranged in a circle (10° diameter), marking the boundary to which participants were instructed to make saccadic responses. At the center of this boundary there was an aperture (5° diameter) inside of which we displayed the random dot motion stimulus. The central stimulus was composed of a white fixation point (0.1° diameter) positioned at the center, and 200 red dots (each 0.1° diameter) with constant velocities of 4 °/s. On each trial we randomly selected a subset of the dots in motion (2%, 10% or 20% of all dots) to move coherently in either the leftward or rightward direction. In the stimulus rotation conditions (HR-S and S), we rotated the on-screen motion axis by either 45° or-45°. In the HR-S condition, this on-screen rotation of motion was congruent with the direction of head roll, such that the motion axis lay approximately along the interocular axis. In all saccadic trials, participants were instructed to make eye movements towards the on-screen 0° (rightward motion) or 180° (leftward motion) directions. Participants were also informed of all block conditions (i.e. head roll, visual stimulus rotation) prior to the start of each block.

A sample trial progression is illustrated in Figure 1B. At the start of each trial, a fixation dot appeared in the center of the circular saccade boundary (fixation period, 500 ms). This fixation period was followed by the visual motion stimulus, displayed within the aperture in the center of the screen along with the fixation point (1500 ms max). Participants were instructed to maintain fixation until they came to a decision about the direction of the coherent motion, and were asked to do so as quickly and as accurately as possible. Depending on the response condition, they either made a saccade along the screen-centered horizontal (left or right) or pressed a button with either their index (left) or middle (right) finger corresponding to the perceived horizontal component of motion. For saccade response trials, participants were instructed to press any button after making a saccade, ending the trial. For button press trials, the decision also ended the trial. Participants were not given feedback about whether their response was correct. There was an inter-trial interval of 500 ms during which the screen was completely black.

Each participant performed four sessions, each consisting of seven, 100-trial blocks for a total of 2800 trials. All 14 conditions (left and right head rolls and stimulus rotations included) were counter-balanced across all participants using a reduced Latin squares method with an initially randomized list of all conditions. To counterbalance potential learning and fatigue effects, participants performed each condition twice; once in an initial sequence determined by the Latin squares method (Shao and Wei, 1992) and a second time in the reverse sequence. Using this method, each condition was uniformly distributed across all blocks.

### Raw signal analysis

3D head orientation was computed offline as the difference (using quaternion rotation based on (Leclercq et al., 2013)) between a reference upright position measured at the start of each experimental session and head positions throughout the trials. Participants were instructed to begin the first block of each experimental session with an upright head position before responding to the verbal head roll instruction.

The eye-in-head orientation was extracted, calibrated and saccades detected using the same techniques as those used by previous work (Blohm and Lefèvre, 2010; Murdison et al., 2013). Briefly, the eye-in-head orientation was extracted after each session from the saved images of the eyes using Iris software (Chronos Vision). This was done using a 9-point grid of calibration dots (10° max eccentricity) with a central fixation point, while the head remained upright on the chin rest. Each participant was fitted with a customized bite-bar to stabilize the Chronos helmet to the head. Eye-in-head orientation was low-pass filtered (autoregressive forward-backward filter, cutoff frequency = 50 Hz) and differentiated twice (weighted central difference algorithm, width= 5 ms). Saccades were detected using an acceleration threshold of 500°/s^2^, as previously done (Blohm and Lefèvre, 2010; Murdison et al., 2013). We defined the eye movement direction as the circular average of horizontal and vertical eye velocity components over the duration of the saccade. For each trial, the head roll measurement was obtained by taking the average head orientation from the motion stimulus onset until the decision time.

### Trial selection

For the main experiment we recorded a total of 19,600 trials from seven participants (2800 trials per participant from four sessions of seven 100-trial blocks each). Of those trials, we removed those that contained a head movement, blink, optokinetic nystagmus or smooth pursuit movement after motion stimulus onset but prior to the decision. Finally, we removed trials on which participants had reaction latencies smaller than 200 ms, as these trials likely represented decisions made preemptively without the use of the visual motion evidence, due to visuomotor processing delays (Thorpe et al., 1996). From the extracted saccades and button presses we determined trial-to-trial directional choices and computed cumulative RT distributions for each rotational condition. For saccades, left or right decisions were classified as saccades whose average direction (based on the entire movement) within a conservative directional window around the screen-centered horizontal direction (0° or 180°) with a width of +/- 75°. Trials with saccades with directions outside these windows were removed from the analysis. Also trials for which the participant failed to respond before the end of the 1500 ms response period were removed from analyses (14% of all trials). Together, these omitted trials comprised 22% of all trials, leaving 15,274 valid trials.

### Behavioral analysis

We quantified task performance using three main behavioral parameters capturing both speed and accuracy aspects of task performance. These parameters were RT (time elapsed between motion stimulus onset and response), percent error (number of valid incorrect trials divided by the total valid correct and incorrect trials; conversely, percent correct = 100%-percent error), and reward rate (sum of the number of correct trials divided by the sum of all correct and incorrect reaction times). From these parameters we computed the cumulative RT distributions for correct and incorrect trials, to which we fit a modified version of the linear approach to threshold with ergodic rate (LATER) model (Carpenter and Williams, 1995).

Because of the short 1500 ms response window some RT distributions were truncated, resulting in LATER-estimated RT distributions that were not necessarily representative of the data. To account for this issue we fit both correct and incorrect trial RT distributions simultaneously using estimated percent correct as a free parameter that scaled each distribution relative to the other correct (representing percent correct or (100%-percent error) at RT = ∞). We also performed all analyses with the empirical percent correct using just the trials within the 1500 ms window and found results qualitatively similar to those based on the estimated percent correct. We performed the fits using a constrained nonlinear method that minimized the sum of squared residuals. These LATER model fits to the cumulative RT distributions revealed the estimated median reaction latency with its *μ* parameter, the approximate slope of the distribution (representing the variability of the distribution) with its *σ* parameter, and the estimated percent correct, each of which we used in behavioral analyses.

We also fit participant and group-level psychometric curves using the Psignifit Toolbox for Matlab (Wichmann and Hill, 2001; Fründ et al., 2011), and fit chronometric data with a scaled logistic function using a nonlinear least squares method. From the psychometric fits we extracted the 75% PSEs and computed the just-noticeable difference (JND) based on the difference threshold, which is a function of the slope and the midpoint percentile for 2AFC tasks *π* (= 75%), described by equations (1) and (2):

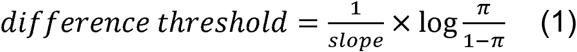

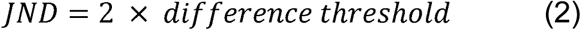

### Reference frame analyses

We then performed a reference frame analysis on the observed behavioral effects for each rotation condition. To do this, we first made predictions for these effect sizes proportional to the complexity of the RFT in each reference frame (Figure 5A), then computed R-squared coefficients for changes (relative to the non-rotated condition) in RT, percent correct and reward rate. These predictions represented RFTs ranging from highly complex (large effect size), intermediately complex (intermediate effect size) or simple (no effect), depending on the angle of coherent motion between input at a given reference frame and the required output, which was left or right for either response type. Because they arose from rotations to retinal input due to ocular torsion during head roll (known as ocular counter-roll), intermediate effect sizes were inversely dependent on one another for eye- and head-centered frames. Instead of choosing arbitrary intermediate effect predictions (e.g. 0.5) we optimized the chosen intermediate predictions for each response effector (eye or hand) and each behavioral parameter (RT, percent error or reward rate). This optimization process chose the intermediate prediction that produced the highest across-participant and across-motion coherence mean R-squared value in our reference frame analysis. For saccades, this optimization yielded eye-centered intermediate predictions of 0.83 (latency), 0.54 (percent error) and 0.59 (reward rate), corresponding to head-centered predictions of 0.17, 0.46 and 0.41 respectively. For button presses, this optimization yielded eye-centered intermediate predictions of 0.55 (latency), 0.55 (percent error) and 0.59 (reward rate), corresponding to head-centered predictions of 0.45, 0.45 and 0.41 respectively.

### Control experiment

We conducted a control experiment in order to account for potential confounds in our data. Seven participants performed four sessions, each consisting of six, 100-trial blocks (2400 trials per participant) for a total of 16,800 trials, of which we removed 17% of trials for reasons previously listed for the main experiment (see *Trial selection*), leaving 13,927 valid trials.

First, we wanted to ensure that any effects we observed in the stimulus rotated condition (S) were due to reference frame transformations and not due to participants only accounting for motion along the screen horizontal, which, in the S condition was decreased by a factor of √2. To compensate, we introduced a new condition in which the speed of the stimulus was increased by a factor of √2 (final speed of 5.7°/s) while the screen stimulus was rotated, called S-spd, depicted in Figure 6A.

Second, we wanted to isolate the variability added to the decision process by the initial sensory estimate of head roll. With this in mind, we introduced a condition only for saccadic responses in which the head, stimulus and saccadic response axis were all rotated congruently, called HR-S-RR, depicted in Figure 6A. Therefore, behavior during this condition could be compared to that during the control condition in order to isolate the variability added by head roll. For completeness we included all of the other conditions in the main experiment, and carried out an identical fully counterbalanced and blocked design.

Finally, we wanted to ensure that truncation of the RT distributions did not play a role in our observations during the main experiment. Participants were again given instructions to “decide as quickly and accurately as possible,” but we allowed them to take up to 5000 ms to decide the direction of coherent motion, rather than 1500 ms. Participants rarely took the full time to reach a decision (0.1% of all trials). Importantly, this task change did not result in any qualitative differences from our main experiment findings.

### Statistical analyses

We performed several n-way ANOVAs (either with 6 or 10-factors, including interaction terms) to account for variance in decision making behavior (across RT, percent error and reward rate) due to coherence level, RFT requirements, participant and motor effector. To correct for statistical sampling error, we also carried out a multiple comparison procedure based on Tukey’s honestly significant difference criterion. We used the 95% confidence intervals estimated using Monte-Carlo simulations (Wichmann and Hill, 2001; Fründ et al., 2011) to compare 75% PSEs and JNDs across RFT conditions in our psychometric analyses.

## Results

We utilized several different rotational conditions to determine the effects of saccade-related (eye-to-head) and button press-related (eye-to-shoulder) reference frame transformations on the performance of a 2AFC perceptual decision task. Using these conditions, we systematically induced different reference frame transformation requirements under which we analyzed the effects on speed (RT), accuracy (percent error), and net performance (reward rate). This approach allowed us to determine both if changing the RFT requirements had any effect on the integration of decision evidence and, if so, if these effects revealed anything about the coordinate frame of the neural circuitry underlying these decisions.

### Head and stimulus rotations induced distinct effects on response times and task performance across conditions

We found that head and stimulus rotations induced different effects on RT and accuracy depending on condition. As shown in Figure 2A for example participant 7, cumulative distributions of RTs showed that, depending on the rotation condition, the estimated median RTs shifted by various amounts relative to the control condition in which the head was upright and the stimulus motion axis was horizontally oriented. We also observed overall increases in RT and decreases in accuracy with task difficulty (20% to 10% to 2% motion coherence), with each condition inducing different effect magnitudes. These effects depended on the response type, suggesting a potential role for the transformation required to convert sensory input into the response frame used for decision making.

**Figure 2:**
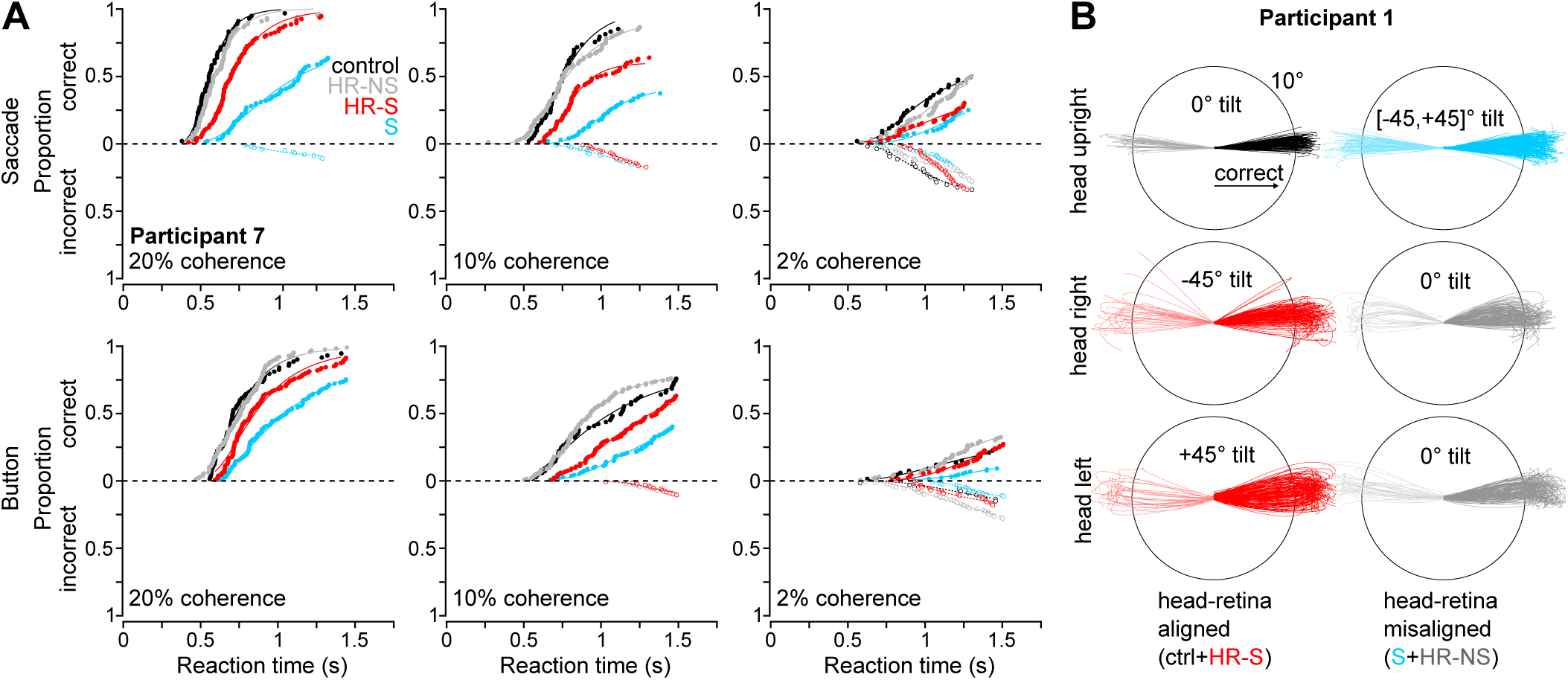
Rotation condition affected RTs and saccade trajectories. (A) Across coherence levels (columns) specific patterns in RTs across rotational conditions (color-coded, see legend) are shown for participant #7. Differences in the order of these RT distributions can be seen when comparing saccade (top row) to button responses (bottom row). (B) Compared to control (upper left panel), saccade trajectories were more variable under rotated conditions.

Although participants were instructed specifically to perform saccades along their perceived screen-horizontal axis, the absence of visual landmarks around the border of the stimulus allowed us to examine how inducing new rotational conditions altered eye movement generation. As can be seen in Figure 2B for example participant 1, changing the rotation condition resulted in more variable saccade trajectories compared to the control condition. Combined with the observed condition-dependent changes to RT and accuracy, these findings suggest that visuomotor transformations systematically affect the neural processes underlying decision making.

### RT and percent correct varied with effector, but there was no speed-accuracy tradeoff

Not only did each rotational condition induce RT and accuracy effects relative to control, but those effects depended on response type. Figure 3 illustrates this phenomenon with psychometric and chronometric functions at the group level. For example, psychometric functions (left column) show that behavior qualitatively differed between conditions depending on whether participants responded with a saccade or button press. Under the HR-NS condition (grey), participants performed similarly to control and reached the overall 75% correct threshold (or 2AFC psychometric PSE) at the lowest motion coherence of any condition (left inset). The JND, which is defined as the ratio representing the units of motion strength (% coherence) required to increase the percent correct (%) by a single unit, reveals that participants also tended to perform with the highest precision in the HR-NS condition (right inset). After the HR-NS condition, the control, HR-S (red) and S (cyan) conditions follow in overall accuracy (PSE) and precision (JND) across motion strengths, with S condition JNDs differing significantly from those of other conditions (*p* < 0.05). Comparing directly to button responses (lower left panel), one can see that this pattern is qualitatively different, with the control condition having the lowest PSE of the conditions (*p* < 0.05), followed by HR-NS, HR-S, and S. The saccadic response chronometric functions (upper right panel), represented across motion coherences and conditions reveal how RT distributions varied with motion coherence and rotational condition. Across all coherence levels, RTs were smallest in the control condition, followed by the nearly identical HR-NS and HR-S conditions, and finally the S condition.

**Figure 3:**
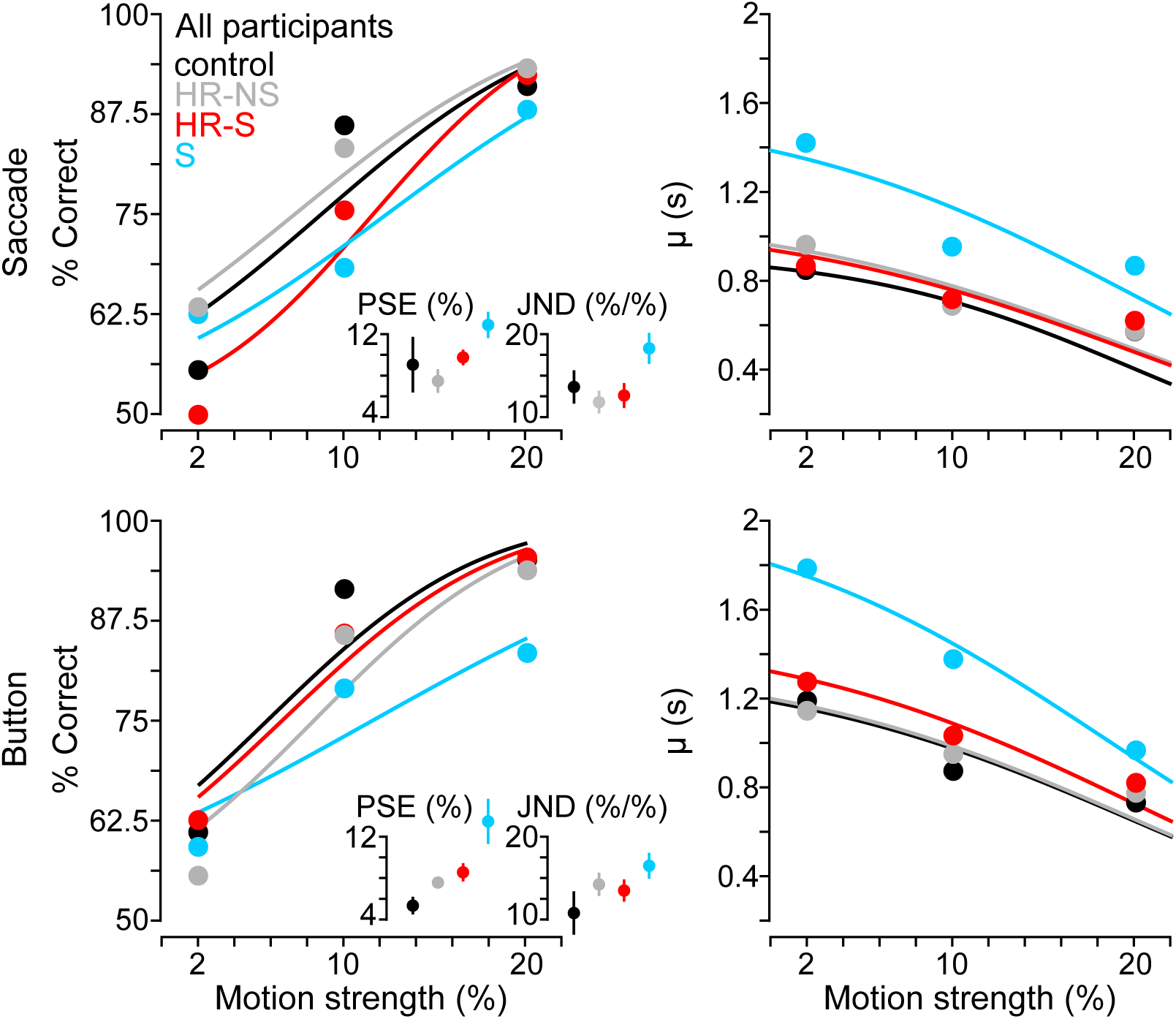
Psychometric and chronometric functions. Group-level psychometric and chronometric functions revealed that speed and accuracy were not traded-off across rotation conditions, as participants were generally less accurate (psychometric functions, left column) and also slower (chronometric functions, right column) under rotated conditions. In the chronometric plots, each point represents the group average of the LATER fit parameter μ approximating the median latency of each condition at each motion strength. Left insets show the point of subjective equality (PSE), which represents the threshold coherence (%) at which participants chose the correct direction 75% of the time for the 2AFC task. On the same scale but with different units (% coherence per % correct), right insets also show the just-noticeable difference (JND), which approximates the amount of variability in the psychometric function using the inverse of the slope around the 75% PSE, scaled for 2AFC tasks. Each of these insets reveals a consistent trend in the median and variability of performance across rotation conditions.

Taken together, these observations suggest that there was an overall degradation of the encoded evidence and no clear speed-accuracy tradeoff (SAT) across rotational conditions. Similarly, button press responses showed a pattern of psychometric (lower left panel) and chronometric (lower right panel) changes suggesting an overall degradation of the encoding of evidence by rotational changes rather than an SAT (Standage et al., 2014b). Additionally, the observed effector specific patterns of performance changes across condition suggest that the reference frame of the motor response played a role in the encoding of evidence.

However, we observed high amounts of inter-participant behavioral variability. We show this variability for changes in RT, percent error and reward rate in Figure 4 across all participants (colored line segments on left axes). We also observed some consistent trends across task difficulty (RT: *F*(2) = 12.73, *p* < 0.01; percent error: *F*(2) = 326.5, *p* < 0.01; reward rate: *F*(2) = 33.54, *p* < 0.01), rotation condition (RT: *F*(3) = 7.78, *p* < 0.01; percent error: *F*(3) = 4.76, *p* < 0.05; reward rate: *F*(2) = 34.25, *p* < 0.01) and effector (reward rate: *F*(1) = 21.58, *p* < 0.01). Note that for reward rate (bottom row), the y-axes are inverted for visualization purposes. On average (inset bars on right axes), participants had longer RTs and had lower reward rates when making decisions under the S condition (cyan bars), when compared to control (multiple comparison *p* < 0.05), HR-NS (grey; multiple comparison *p* < 0.05), and HR-S (red; multiple comparison *p* < 0.05) conditions. Importantly, we did not see an SAT, as reward rate also decreased (bottom row) with increases in both RT and percent error. These behavioral changes were, however, consistent with a degradation of evidence encoding such that the task was more difficult under rotated conditions, and that difficulty increased with RFT complexity. We observed participant-specific differences in RT between effectors (interaction effect, *F*(6) = 4.93, *p* < 0.01) and between RFT condition (interaction effect, *F*(18) = 3.03, *p* < 0.01). For example, one can see differences between saccade and button responses for participant #5 (blue traces) or for participant #3 (yellow traces) across each effector and coherence level. This trend suggests that the noise added to the evidence encoding not only changed with effector, but also with rotational condition, in agreement with the observed changes to psychometric and chronometric functions. We next used a reference frame approach to determine the source of this additive noise in the decision process.

**Figure 4:**
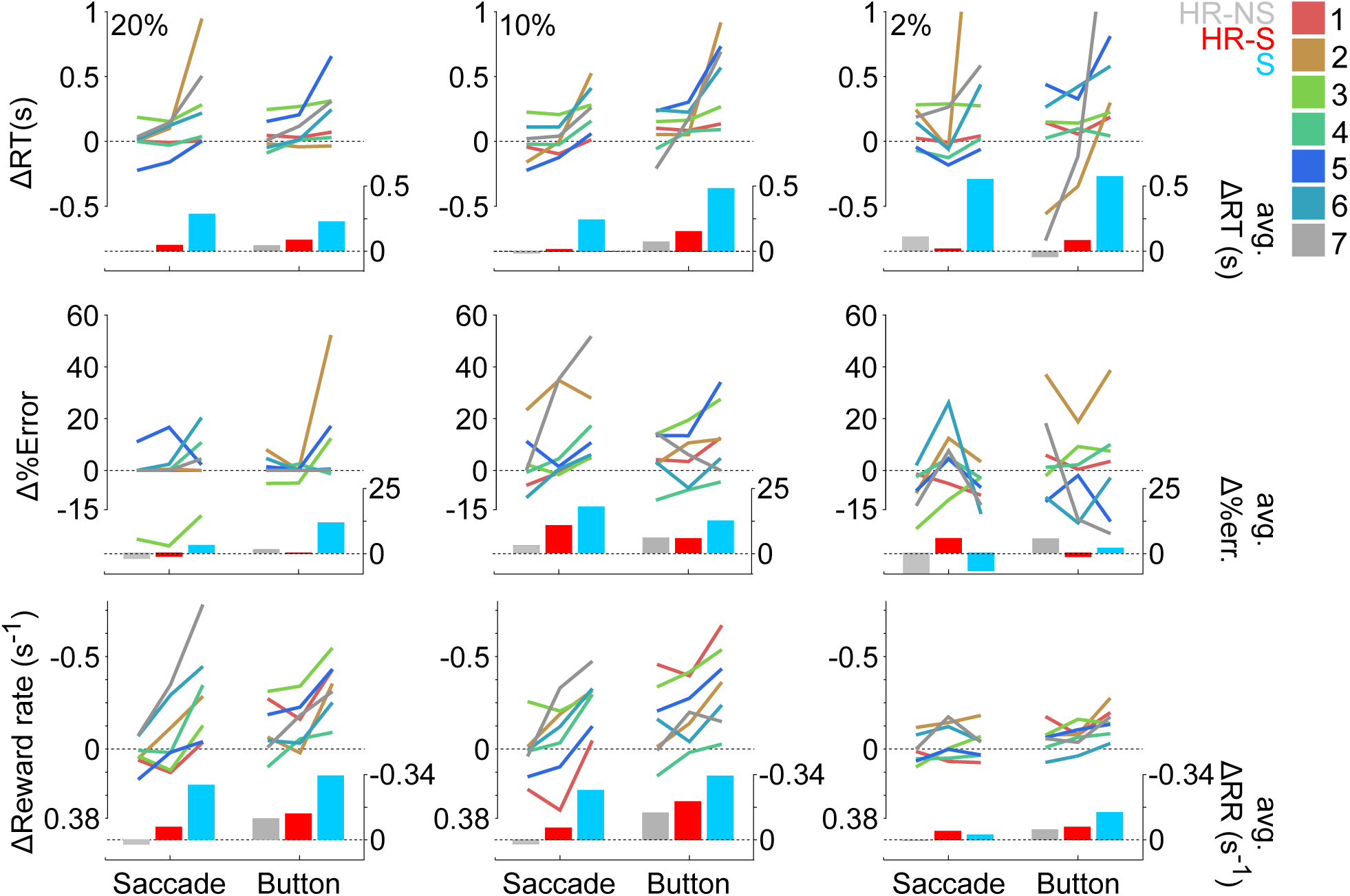
Variability of rotational effects on performance across participants. Changes in latency (top row), percent error (middle row) and reward rate (bottom row) across coherence level (columns), with left axes representing scale for single participant changes (colored line segments, see legend for participant numbers) and right axes representing group-level average changes across rotation conditions (color-coded bars). Each vertex of the line segments represents one rotation condition, in line with the colored bars at the bottom. Note that for direct comparisons with latency and percent error changes we inverted the vertical axes for reward rate changes.

### Reference frame analysis

To quantify this inter-participant variability, we interpreted the effects using predictions from stochastic reference frame transformations (Alikhanian et al., 2015). We did this under the assumption that the motion information used in the decision was impaired to an extent that was proportional to the complexity of the required visuomotor rotation. Using this approach, we predicted the size of each effect, relative to control, according to the required rotation for a correct effector-centered response in each condition, which we illustrate in Figure 5A. For example, consider the eye-centered prediction for the condition in which both the head and the screen were rotated and a saccadic response was required (HR-S; middle cell, top row, top grid, panel A): in order to correctly interpret the spatial motion direction using eye-centered information, the brain must rotate the retinal vector (which points along its horizontal; for visualization see Figure 1A) by the head roll magnitude to generate a screen-centered horizontal saccade. This requirement differs for the condition in which the head, but not the stimulus, was rotated (HR-NS). Because the retinal vector was rotated solely by head roll and ocular counter-roll, and the eyes are also rotated along with the head, the brain only needed to account for ocular counter-roll when transforming the retinal vector into a screen-horizontal saccade. Therefore, in the eye-centered case, we predicted a large stochastic effect for HR-S (black shading) due to head roll and an intermediate effect for HR-NS (grey shading) due to only ocular counter-roll. In this way, we made predictions for each effector and for each reference frame (eye, head and shoulder).

**Figure 5:**
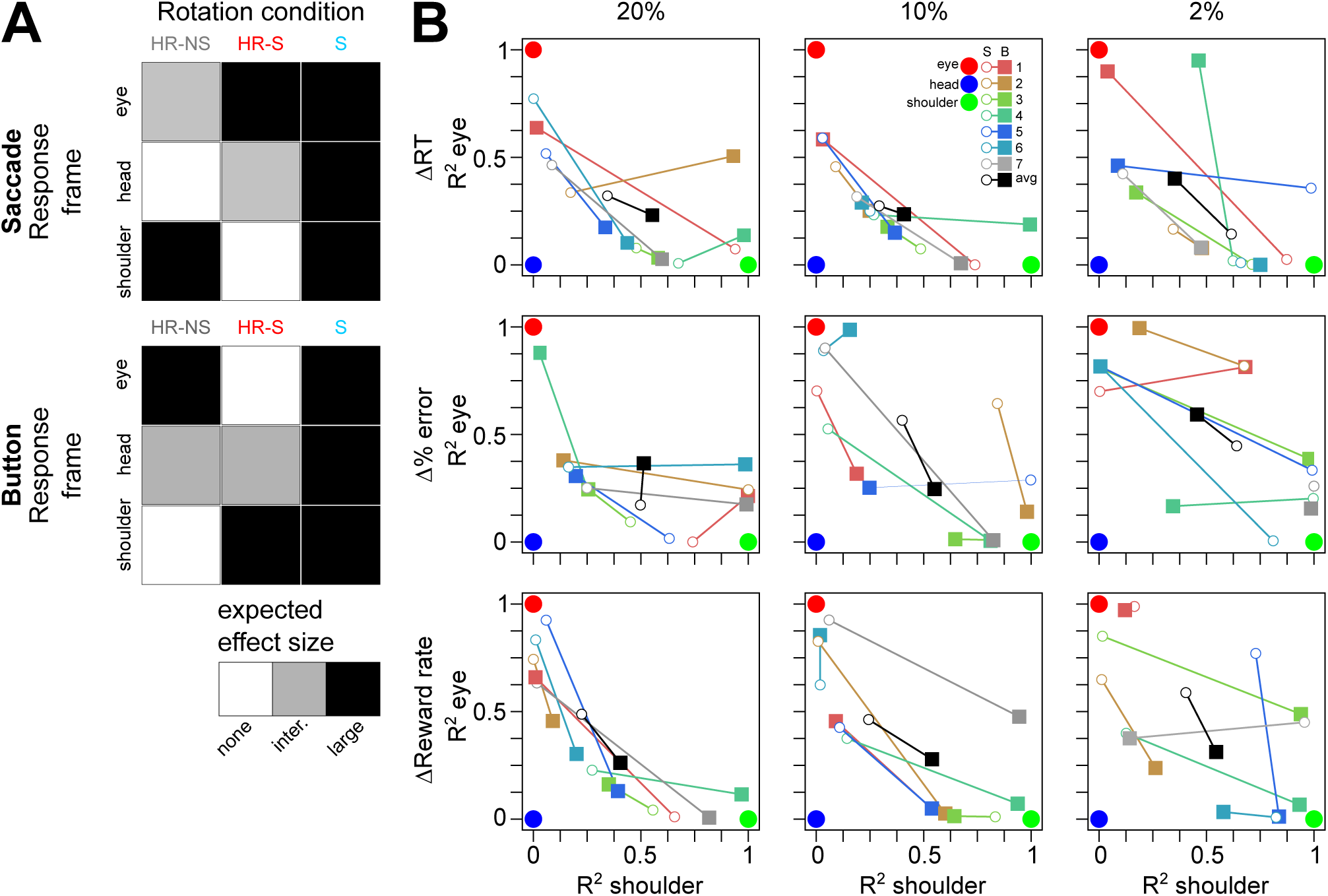
Reference frame predictions and analysis. (A) Effector specific reference frame prediction matrices. Each cell represents a specific reference frame and the predicted effect size for the corresponding rotation condition. For example, if motion evidence were coded according to an eye-centered reference frame, for the condition in which only the motion stimulus were rotated (condition S) we would expect a large (black shading) reference frame transformation-induced stochastic effect on the coded evidence signal in both saccade and button response conditions. (A) Participant R-squared coefficients for correlation analysis between prediction matrices in panel (A) and observed changes in latency (top row), percent error (middle row), and reward rate (bottom row), across coherence levels (columns). Participant color code is the same as in previous figures, and black symbols represent across- participant means. Open circles and filled squares represent R-squared coefficients for saccade responses and for button responses, respectively. Pure eye- (red), head- (blue), and shoulder-centered (green) reference frame predictions are represented with large filled circles.

Using these predictions, we computed the R-squared coefficients for each behavioral parameter (RT, percent error and reward rate), each participant, each effector and each motion coherence. These are depicted in Figure 4B along with the predictions for a purely eye-centered (red dot), head-centered (blue dot) and shoulder-centered (green dot) codings. Each R-square coefficient is color-coded according to participant and represented by a symbol depending on response type (saccades: open disk, button: filled square). Across both RT and percent error at 20% coherence, the R-square coefficients suggest that evidence was being encoded according to a continuum of reference frames between eye and shoulder, with a strong head-centered component in some cases (e.g. button press responses of participant 5).

The transformation-related effect was also dependent on the strength of the stimulus, indicating that the addition of variability to the encoded evidence depended on the initial strength of visual motion. For example, while there is a clear organization of R-square coefficients for the 20% and 10% motion coherence conditions for changes in latency along an eye-head-shoulder continuum (Fig 5B, upper left and middle panels), this continuum becomes less clear when the stimulus strength is decreased at 2% motion coherence (Fig 5B, upper right panel).

With this analysis, we quantified the effector specific component that we initially observed in the psychometric and chronometric functions (Figure 3). This component was strongest when considering reward rate (bottom row of Figure 5B). Across motion coherence, group reward rate averages (black symbols) indicated that evidence leading to saccadic responses was more eye-centered while evidence leading to button responses was more shoulder-centered. This trend suggests that the neural circuitry encoding decision evidence is tied to the motor plan for the upcoming movement. Additionally, this mixture of eye- and shoulder-centered components indicates that there could be some concomitant evidence coding by eye- and shoulder-related areas during integration, regardless of eventual motor effector.

### Control experiment

Our main experiment had two important limitations: (1) in the stimulus-rotated condition S we could not definitively rule out the influence of decreased motion energy along the screen horizontal during the integration of motion evidence, and (2) we could not isolate the effects of only head roll on the decision process. To address these limitations, we re-ran the experiment with a new group of participants with two added conditions: (1) screen rotation with a proportional increase in the speed of the stimulus to compensate for the loss of horizontal motion energy in the initial S condition (S-speed, green) and (2) head roll with rotation of the screen stimulus and saccadic responses rotated along the motion axis, and not screen horizontal (HR-S-RR, purple), depicted in Figure 6A.

**Figure 6:**
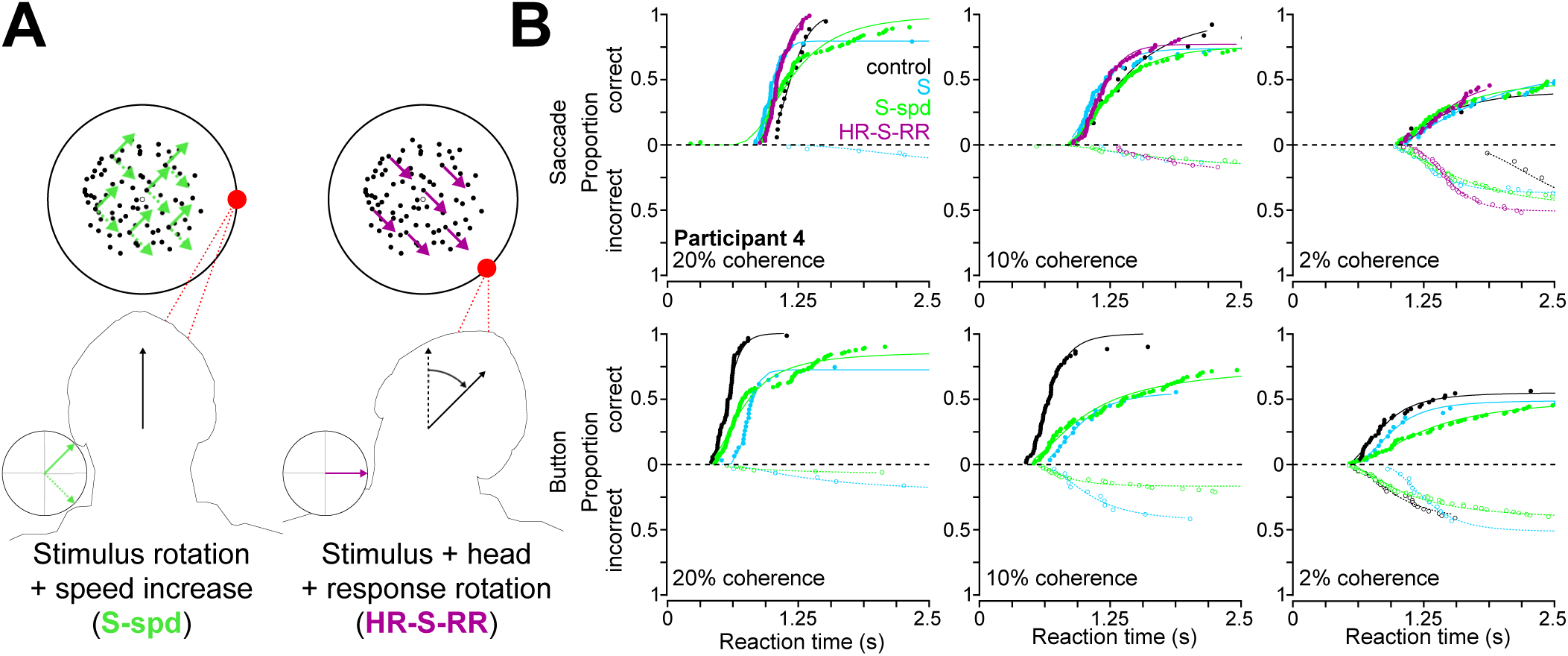
Control experiment conditions and example RT distributions. (A) Representation of added rotational conditions in which the speed of the motion stimulus was increased to compensate for the loss of horizontal motion in the S condition (S-spd, left) and in which the saccadic responses were also rotated to match the rotated motion axis, thus isolating the effect of head roll in the transformation (HR-S- RR, right). (B) Exemplar participant RT distributions for saccades (top row) and button responses (bottom row), showing control (black), S (cyan), S-spd (green) and HR-S-RR (purple) conditions across coherence levels (columns).

Importantly, this experiment produced similar statistical RT, accuracy and reward rate effects as the main experiment for the repeated RFT conditions across task difficulty, motor effector, rotation condition and participant. Shown in Figure 6B, the cumulative RTs for participant 4 show that both the S-spd and HR-S-RR conditions each produced behaviors similar to their conditional counterparts (note that this is a different participant 4 than in the main experiment). We detected no differences in RT, percent error or reward rate due to the RFT between S-spd and S or between HR-S-RR and HR-S, but found one significant RFT effect between control and HR-S-RR for only percent error (*F*(1) = 9.10, *p* = 0.03, for RT and reward rate all *p* > 0.05). These findings indicate that (1) there was no detectable behavioral effect of the decrease in horizontal motion energy during the S condition in the main experiment, thus validating our initial findings, and (2) the behavioral effects we observed under head roll conditions resulted from the transformation itself and not from a noisy initial sensory estimate of head roll.

## Discussion

### Summary of findings

The goal of this study was to determine the influence of stochastic reference frame transformations on decision making. We designed a paradigm in which 7 participants performed a 2AFC motion direction discrimination task under control conditions (head upright, stimulus motion along the screen horizontal) or under one of several rotation conditions in which the head and/or stimulus were rotated. Combining rotation conditions with saccadic and button responses allowed us to behaviorally quantify eye-, head- and shoulder-centered effects.

We made predictions for the influence of RFTs on speed (RT), accuracy (percent error/correct) and overall performance (reward rate). We found (1) that stochastic reference frame transformations impair decision making, leading to slower, less accurate decisions, (2) that this stochasticity is added in a manner consistent with a mixed eye-head-shoulder representation of evidence, and (3) that within this continuum there is an effector specific component, with saccadic responses more closely resembling eye-centered predictions and button responses more closely resembling shoulder-centered predictions. Our findings are consistent with the hypothesis that perceptual decision making and visuomotor reference frame transformations occur within the same neural circuitry (Dorris et al., 1997; Gold and Shadlen, 2000), and as such are consistent with the affordance competition hypothesis of embodied decision making, which predicts that motor planning for perceptual decision making occurs in parallel between networks coding for multiple potential actions (for reviews see Cisek 2007; Cisek and Pastor-Bernier 2014).

Although both evidence integration and motor preparation are often necessary for choice behavior, it is often difficult to distinguish between the contributions of each using standard perceptual tasks. Previous efforts to do so include using delays between stimulus viewing and motor response (Shadlen and Newsome, 2001; Sommer and Wurtz, 2001; Lemus et al., 2007), limiting stimulus viewing time (Bergen and Julesz, 1983; Ratcliff and Rouder, 2000; Bodelón et al., 2007; Kiani et al., 2008) and even “compelling” the movement by informing the perceptual system ahead of time about the target characteristics (Salinas et al., 2014). At the neural level, perceptual and motor processes both occur in sensorimotor association areas (Munoz and Wurtz, 1995; Dorris et al., 1997; Horwitz and Newsome, 1999; Shadlen and Newsome, 2001; Hernández et al., 2010; Costello et al., 2013; Mante et al., 2013). Not only are our findings consistent with these neurophysiological principles, but we have also now quantified this inseparability for the first time within an RFT framework.

### Open questions

We found that transformation-induced stochasticity impairs decision making. Given that psychophysical thresholds are systematically lowered by the added RFT noise, the simplest explanation points to a degradation of the encoding of motion evidence most likely in the middle temporal (MT) or medial superior temporal (MST) areas (Albright, 1984; Britten et al., 1992, 1993, 1996; Salzman et al., 1992; Inaba et al., 2007). MT and MST are highly interconnected areas that serve as the interface between retinal motion signals and the rest of the visuomotor pathways (Ungerleider and Desimone 1986; Komatsu and Wurtz 1988; Newsome et al. 1988; Ilg and Thier 2003; Inaba et al. 2011; for review see Krauzlis 2004), and exhibit gain modulation and receptive field shifts (Chukoskie and Movshon, 2009; Fujiwara et al., 2011; Inaba et al., 2011) mechanistically consistent with carrying out 3D visuomotor transformations (Blohm and Crawford, 2007; Blohm et al., 2009; Blohm and Lefèvre, 2010; Blohm, 2012; Murdison et al., 2015). If these areas indeed provide the neural substrate for the addition of variability to visual motion signals via RFTs, then gain modulation for RFTs itself could be a stochastic process – a question that should be investigated in future electrophysiological and modeling work.

Our findings suggest that the encoding of evidence was shared by effectors. However, the contents of that signal differed between participants – something that might be explained by inter-participant differences in the variability of the evidence integration. RFT stochasticity added to the integration process could result in a less reliable population ‘readout’ of the current estimate of stimulus motion by downstream areas, resulting in more variable RT distributions with shallower slopes (Carpenter and Williams, 1995). Although we did not see any clear indications of this on LATER model slopes (a; see Methods), differences in how these population responses are decoded by structures closer to the motor output such as the superior colliculus (SC) (Munoz and Wurtz, 1995; Dorris et al., 1997; Horwitz and Newsome, 1999; Sommer and Wurtz, 2001) or primary motor cortex (M1) (Riehle and Requin, 1989; Crammond and Kalaska, 1996, 2000) could potentially explain some of the inter-participant variability we observed in RT, percent error and reward rate correlations.

### Potential mechanism and underlying neural circuitry

Our findings are consistent with the hypothesis that the encoding of motion evidence is degraded by RFTs; however, this was not the only possible way that RFTs could have affected decision making. For example, changes in background noise could have modulated the dynamics of circuitry integrating evidence (Furman and Wang, 2008; Roxin and Ledberg, 2008; Standage et al., 2013, 2014a, 2014b), consistent with recent data (Heitz and Schall, 2012). If so, SAT would have been observed (Standage et al., 2014b).

The finding that the impairment of performance relied partially on the response type implies the existence of two partially distinct perceptual decision making networks between behavioral effectors, as previously theorized (Dean et al., 2011; Madlon-Kay et al., 2013). In the lateral intraparietal area (LIP) and the parietal reach region (PRR), which lies along the medial bank of the intraparietal sulcus (IPS), population-level neural activity has been shown to reflect an effector-nonspecific movement signal until a monkey makes a decision regarding which effector to use, at which point PRR activity is associated with a reach (Cui and Andersen, 2007; Yttri et al., 2014; Wong et al., 2016) or LIP activity is associated with a saccade (Cui and Andersen, 2007; Wong et al., 2016). To accomplish this, recent electrophysiological findings (Wong et al., 2016) indicate that there are ensembles of neurons on both the medial and lateral banks of the IPS that are active during the decision process. Specifically, Wong and colleagues (2016) found an ensemble of neurons that predict the upcoming decision, independent of effector specific region, that coherently spike prior to effector specific local ensembles in each bank (Wong et al., 2016), consistent with previous findings (Cui and Andersen, 2007; Yttri et al., 2014). These partially distinct neural ensembles could therefore give rise to the mixture of reference frames our perceptual findings imply should be present in the neural integration of motion evidence. Of course, this explanation does not preclude perceptual and motor contributions from other effector nonspecific areas such as the prefrontal cortex (Madlon-Kay et al., 2013) or from other effector specific areas whose activities are believed to implement a decision variable such as FEF (Hanes and Schall, 1996; Gold and Shadlen, 2000, 2003; Sommer and Wurtz, 2001) or the dorsal premotor cortex (Crammond and Kalaska, 1996, 2000; Cisek and Kalaska, 2002, 2005), or downstream in SC (Munoz and Wurtz, 1995; Dorris et al., 1997; Sommer and Wurtz, 2001; White et al., 2013) or M1 (Riehle and Requin, 1989; Crammond and Kalaska, 1996, 2000). The precise role that RFT stochasticity plays within such a distributed perceptual decision network, especially with several anatomically distinct sensorimotor association areas with different information flow characteristics and latencies is unclear (Siegel et al., 2015). Furthermore, within these areas, it is also unclear how local neural population codes vary with body and spatial geometry during visuomotor decisions. These are questions that should be further investigated psychophysically and electrophysiologically.

Our findings have implications for studies involving the integration of evidence for movement, whether used for perceptual decision making or motor preparation. First, we found that RFT stochasticity affects the encoding of evidence for perceptual decision making, bringing to light the requirement for controlling the visuomotor geometry during perceptual tasks. Second, the finding that this added variability was partially effector specific could explain some variability between psychophysical performance when the perceptual task is identical with the exception of the motor response (Palmer et al., 2005).

The influence of RFT stochasticity on perceptual decision making is consistent with previous findings in visuomotor tasks (Sober and Sabes, 2003, 2005; Schlicht and Schrater, 2007; McGuire and Sabes, 2009; Burns and Blohm, 2010; Burns et al., 2011), suggesting that it represents a generalized phenomenon wherever RFTs can be found throughout the perceptual and motor systems. Whether this phenomenon can be further extended to processes requiring a higher degree of cognitive involvement such as strategic decision making or memory storage and retrieval remain open questions.

## Acknowledgments

The authors want to thank the participants for their kind participation in the study and Ashley Clare Parr for her helpful comments on the manuscript. This work was supported by NSERC (Canada), CFI (Canada), the Botterell Fund (Queen’s University, Kingston, ON, Canada) and ORF (Canada). TSM was partly supported by DAAD (Germany).

